# Decay and Solid-Liquid Partitioning of Mpox and Vaccinia Viruses in Primary Influent and Settled Solids to Guide Wastewater-Based Epidemiology Practices

**DOI:** 10.1101/2025.02.11.633197

**Authors:** Jacob R. Phaneuf, Gyuhyon Cha, Janet K. Hatt, Konstantinos T. Konstantinidis, Katherine E. Graham

## Abstract

Wastewater-based epidemiology (WBE) has proven to be a powerful tool for tracking the spread of viral pathogens, such as SARS-CoV-2, but as WBE has expanded to include new pathogens, such as mpox virus, more data is needed to guide practitioners on how to design WBE campaigns. Here, we investigated the decay rates of heat-inactivated mpox (HI-MPXV) and attenuated vaccinia virus (VV) in primary influent and settled solids collected from a local POTW at 4°C, 22°C, or 35°C using digital PCR. Subsequently, we studied the solid-liquid partitioning of the viruses in primary influent. Over the 30-day study period, we observed no significant difference in log-linear decay rates between viruses (p=0.5258), with significantly higher decay rates in primary influent (0.109-0.144/day) compared to settled solids (0.019-0.040/day) at both 22°C (p=0.0030) and 35°C (p=0.0166). Furthermore, as part of the partitioning experiment, we found that HI-MPXV and VV adsorb to the solids fraction of primary influent at higher intensities than previously studied enveloped viruses (K_F_ = 1,000-31,800 mL/g, n = 1.01-1.41). Likewise, it was determined in the partitioning experiment that a concentration of greater than 10^3^ gc/mL in raw influent was needed for the viable quantification of mpox and vaccinia viruses in the clarified liquid fraction of raw primarily influent. Our study provides essential insights into informative sample collection and storage conditions for the analysis of wastewater and for transport modeling studies. Due to the slow decay observed in settled solids at all tested temperatures in the persistence experiment, this matrix may be most suitable for retrospective analyses of community infection of the mpox virus.

## 1. Introduction

Wastewater-based epidemiology (WBE) has proven to be a powerful tool for community health monitoring by providing insights into infection trends through the analysis of wastewater samples for targeted pathogens. By quantifying specific viral genetic markers, WBE enables public health agencies to assess disease trends within populations served by municipal treatment facilities from which the initial sample is collected. This approach can provide near real-time outbreak data that supplement individual-level disease surveillance practices. During public health crises, such as the COVID-19 pandemic, WBE has proven to be particularly valuable for accurately gauging community infection trends even at the national scale; for instance, through the CDC’s National Wastewater Surveillance System (“CDC: National Wastewater Surveillance System,” 2023). While this practice can enhance our understanding of population health dynamics and inform public health policies, it is necessary to understand the physicochemical effects of wastewater on pathogens, as well as optimized methods for sample acquisition and storage, for further implementation to other pathogens.

Beyond SARS-CoV-2 (Wolfe et al., 2022, 2021; Yu et al., 2022), another virus of significant public health concern that has been successfully detected and monitored in wastewater is mpox virus (formerly monkeypox virus), arising from the 2022 global outbreak and now formally considered an international public health concern (Adams et al., 2024; Sharkey et al., 2023; Wannigama et al., 2023). Historically endemic to Central and West Africa, mpox virus is an enveloped, zoonotic virus with a large double-stranded DNA genome ∼197,205 bp in length (Karagoz et al., 2023) and a brick-shaped morphology of 200 nm by 250 nm, making it an order of magnitude larger in genome length and cell size than previously monitored viruses (Lu et al., 2023). Mpox virus infections can cause skin lesions, high fever, and severe muscle aches, among other symptoms(“World Health Organization: Mpox,” 2024). It is spread through close or direct contact with contaminated materials or other infected individuals. As of 2024, thousands of cases have been reported in Europe, South America, and the United States, prompting the director of the World Health Organization (WHO) to acknowledge the international outbreak of mpox a public health emergency of international concern (“World Health Organization: News,” 2024). The international resurgence of mpox infections highlight the need for robust surveillance, preventive measures, and public health interventions to mitigate its impact and prevent future outbreaks, which could be informed by WBE.

While WBE has the potential to provide insight into mpox community infectivity and guide public health decision-making to combat the escalation of current global outbreaks, the analyses associated with it are limited by the lack of research and understanding of the fate and persistence of poxviruses in wastewater. A systematic review (Silverman and Boehm, 2021) found only four decay constants of orthopoxviruses from two different studies. However, these studies studied only one species of orthopoxvirus (vaccinia virus) in seawater (Magnusson et al., 1966) and stormwater matrices (Essbauer et al., 2007), stressing a critical knowledge gap in the generalizability of these values for mpox virus in wastewater specifically. This lack of targeted research emphasizes the urgent need to examine how orthopoxviruses, such as mpox virus, behave across different wastewater matrices to inform WBE monitoring (Maal-Bared et al., 2022). For instance, understanding both the decay rates and solid-liquid partitioning behavior of mpox DNA markers in wastewater can influence sampling strategies and the interpretation of data. As mpox continues to pose a threat to global public health, such research will be essential for developing effective mitigation strategies and ensuring that wastewater monitoring can accurately reflect community infection levels.

In this study, we posed the following research questions: 1) how do decay rates vary for two (attenuated) orthopoxviruses in samples of primary influent and settled solids at varying temperatures? and 2) how do the two orthopoxviruses partition between wastewater liquids and solids? To answer these questions, we constructed and spiked wastewater mesocosms with viruses to investigate how virus type, matrix, and temperature affect the decay rates of poxviruses. Primary influent and settled solids were collected from POTWs, aliquoted into different mesocosms, and spiked with orthopoxviruses. Mesocosms were shielded from light, stored at different temperatures, and subsampled over the 30-day study period. We also conducted a solid-liquid partitioning experiment to better understand the adsorption capacity and intensity of these two viruses. Ultimately, the data we provide here will be useful for researchers modeling the fate of viruses in sewer systems, as well as practitioners of WBE who are deploying orthopoxvirus surveillance programs.

## 2. Methods

Stocks of attenuated or heat-inactivated orthopoxviruses (BSL-2) for spiking into wastewater samples were obtained from BEI Resources or the American Type Culture Collection (ATCC): heat-inactivated mpox virus (obtained through BEI Resources, NIAID, NIH: Monkeypox Virus, hMPXV/USA/MA001/2022, Lineage B.1, Clade IIb, cat. no. NR-58711) (referred to as HI-MPXV throughout) and vaccinia virus strain Modified vaccinia virus Ankara (MVA) (cat. no. VR-1508, referred to as VV throughout). By including both HI-MPXV and VV, we can acquire more data on the behavior of these two viruses in wastewater and better understand their generalizability. Viruses were thawed on ice and diluted in filter-sterilized 1X PBS to create separate virus working stocks.

### 2.1. Methods evaluation experiment

Primary settled solids (1 L) were collected from two POTWs in Atlanta, GA, on March 22, 2023, and stored on ice during transport to the lab. From POTW-A, it was a grab sample and from POTW-B, it was a 24-hour composite sample. One 24-hour composite primary influent sample (post-grit screening) was also collected from POTW-A. In the lab, pH and total suspended solids were measured for each sample. Wastewater samples were split into three replicates for spiking with two surrogate viruses: (1) heat-inactivated monkeypox virus (HI-MPXV) and (2) attenuated vaccinia virus (VV) (ATCC cat. no. VR-1508). A background sample was also obtained which was not spiked to account for any endogenous markers in the collected samples (all below the limit of detection). Primary settled solids samples from two different POTWs were spiked with stocks of virus solutions to a final concentration of ∼10^5^ gene copies (gc) per gram of wet solids (HI-MPXV) or ∼10^6^ gc per gram of wet solids (VV). The influent sample was spiked with ∼10^5^ gc (HI-MPXV) or ∼10^6^ gc (VV) per milliliter. The “no spike” group did not receive any additional biological materials spiked into the samples. All samples were inverted to mix and left to equilibrate for 30 minutes (min) on ice prior to additional sample processing.

Primary settled solids samples were concentrated within 24 hours of collection and spiking using either a dewatering method (Loeb et al., 2020) or a protocol adapted for Nanotrap beads from sludge (Ceres Nanosciences Inc., Manasas, VA). Raw primary settled solids, which served as a molecular process control (MPC, i.e., no concentration step for this sample), was also retained for spiking and extractions and preserved in lysis buffer at 4°C prior to nucleic acid extraction. Influent was concentrated using either a membrane filtration method with MgCl_2_ addition (Ahmed et al., 2020b) or Nanotrap magnetic bead method (Karthikeyan et al., 2022). Concentrates were stored in lysis buffer at 4°C prior to nucleic acid extraction within two days of collection. Raw influent, serving as a molecular process control (MPC, i.e., no concentration step for this sample), was also retained for nucleic acid extractions. More details on the methods can be found in the Supplementary Material (SM).

In addition to the comparison of the above concentration methods, we also compared extraction methods. In addition to concentrates, separate raw primary influent and settled solids samples were spiked with either HI-MPXV or VV and directly extracted using the following methods, to serve as molecular process controls (MPCs). In this way, the unconcentrated wastewater samples can provide estimates of recovery of viruses without losses associated with concentration steps. Primary settled solids concentrates were extracted using either the Qiagen DNeasy PowerSoil Pro kit (cat. no. 47014) or Qiagen DNeasy PowerMax Soil kit (cat. no. 12988-10, for the dewatered primary settled solids samples only). Both sets of extractions were completed using the manufacturer’s protocol, utilizing bead beating in the provided tubes. Influent samples were extracted using either the Qiagen AllPrep PowerViral DNA/RNA kit or Omega Biotek MagBead Viral DNA/RNA kit following the manufacturer’s instructions (final elution volume = 100µL). Blank extraction negative controls were created by substituting 1X filter-sterilized PBS for samples and extracted using each method above.

### 2.2. Persistence Experiment

An additional 24-hour composite primary influent and primary settled solids sample (1 L each) were collected from POTW-A in Atlanta, Georgia, on February 9, 2024, and stored on ice during transport to the lab. The facility reported a flow rate of 49.65 million gallons per day (MGD), total suspended solids (TSS) concentration of 366 mg/L, carbonaceous biological oxygen demand (CBOD_5_) of 216 mg/L, and total phosphorus (as P) concentration of 16.2 mg/L at the time of sample collection. To measure percent solids, a subsample of the primary settled solids sample was heated on a pre-weighed aluminum dish at 105°C for 24 hours and mass was measured both before and after the heating in accordance to a previously published method (Graham et al., 2021).

Samples were processed within 24 hours of collection. Both the primary influent and settled solids samples were thoroughly mixed prior to mesocosm set-up by inverting each 1 L sampling bottle thrice and vortexing individual mesocosms for 30 seconds following sample addition. A set of mesocosms using DNase/RNase Free molecular-grade water was used as a third condition to understand the effect of matrix on decay rates. For the primary influent and DNase/RNase Free molecular-grade water mesocosms, 1.5 mL was aliquoted into 2 mL DNase/RNase-free tubes. For the primary settled solids mesocosms, 1.5 g was aliquoted into 5 mL DNase/RNase-free centrifuge tubes. Each matrix mesocosm was produced in triplicate for each of the three temperatures tested: 4°C, 22°C, and 35°C, and for two different virus spikes, HI-MPXV and VV (n = 54). Temperatures were chosen based on comparisons to published manuscripts and to represent a variety of sewer and storage conditions. For the HI-MPXV spike, 30 µL of a 1.25 x 10^3^ gc/µL working solution was added to yield a final concentration of 2.50 x 10^4^ gc/mL_influent_ or 2.50 x 10^4^ gc/g_raw solids_ in each mesocosm. For the VV spike, 6 µL of a 6.25 x 10^4^ gc/µL working solution was added to yield a final concentration of 2.50 x 10^5^ gc/mL_influent_ or 2.50 x 10^5^ gc/g_raw solids_ in each mesocosm. Additionally, primary influent and settled solids samples were taken prior to spiking to assess any potential background signal of viral DNA presence and stored at 22°C (n = 2). Three different no spike DNase/RNase Free molecular-grade water mesocosms were produced as negative spiking controls during sample processing, one for each temperature (n = 3). Mesocosms were placed into opaque boxes to shield them from ambient light and placed into their respective constant temperature rooms.

On Days 0, 4, 8, 14, and 30, each mesocosm and process controls were subsampled for immediate DNA extraction using either the Qiagen AllPrep PowerViral extraction kit (with beta-mercaptoethanol and homogenization) for the primary influent and DNase/RNase Free molecular-grade water mesocosms, while the Qiagen DNeasy PowerSoil Pro extraction kit was used for the primary settled solids mesocosms. For the primary influent and DNase/RNase Free molecular-grade water samples, each mesocosm was vortexed for 10 seconds and 200 µL was collected for DNA extraction. Extractions using the PowerViral kit were done using the manufacturer’s protocol using beta-mercaptoethanol and bead-beating for 10 min at room temperature on a FastPrep-24 5G bead-beater (MP Biomedicals). For the primary settled solids samples, each mesocosm was vortexed for 10 seconds and approximately 250 mg of solids was aliquoted into pre-weighed 2 mL tubes. Actual mass measurements taken for each sample can be viewed and downloaded from the link found in the Data Availability section. Each subsample tube was centrifuged at 4200 rpm for 40 min at 4°C (Roldan-Hernandez et al., 2022) After centrifugation, the liquid supernatant was discarded, the remaining solids mass was measured, and DNA was extracted using the PowerSoil kit manufacturer’s protocol and bead-beating for 10 min at room temperature on a FastPrep-24 5G bead-beater (MP Biomedicals). Blanks were included for both kits with each extraction batch (n = 10). Nucleic acid extracts were stored at -80°C until assayed. Extracts were assayed for HI-MPXV and VV by dPCR within 3 months of extraction, following the same assay and conditions as detailed in the previous section.

Due to low concentrations in the HI-MPXV solids, six wells were dedicated to each sample and analyzed using the HyperWell setting within the Qiacuity software and accounted for in the dimensional analysis. All other samples were assayed in single wells.

### 2.3. Partitioning Experiment

A third 24-hour composite primary influent was collected from POTW-A and stored on ice during transport to the lab. The facility reported a flowrate of 36.7 MGD, TSS concentration of 124 mg/L, CBOD_5_ of 101 mg/L, and total phosphorus (as P) concentration of 5.9 mg/L. The primary influent was inverted 3 times to mix. After, 40mL of influent was aliquoted into a 50mL conical tube. HI-MPXV was obtained from ZeptoMetrix (Strain: USA/MA001/2022, SKU: 0810657CFHI) and diluted to a working solution of approximately 2.50 x 10^4^ gc/µL using 1x filter-sterilized PBS. Purification of both viruses was performed by passing the working stock through an Amicon Ultracentrifugal 100 kDa filter (cat. no. UFC5100) following the manufacturer’s protocol. The purified viruses were then resuspended in filter-sterilized 1x PBS and sonicated for 90 seconds at a frequency of 40 kHz using a Fisher Scientific Ultrasonic Cleaner (SKU:19076) to dissociate aggregates, based on a previously published study (Galasso and Sharp, 1962), and the final concentration was measured by dPCR.

For HI-MPXV spikes, four different final concentrations were produced in duplicate (n = 8). Tubes were spiked with the working solution to achieve a final concentration of approximately 2.5 x 10^3^, 2.5 x 10^4^, 2.5 x 10^5^, and 2.5 x 10^6^ gc/mL, respectively. For VV spikes, four different final concentrations were produced in duplicate (n = 8). A 6.25 x 10^4^ gc/µL working solution was added to yield final concentrations of approximately 1.25 x 10^3^ gc/mL, 1.25 x 10^4^ gc/mL, 1.25 x 10^5^ gc/mL, and 1.25 x 10^6^ gc/mL. Additionally, no spike primary influent and DNase/RNase Free molecular-grade water 40mL samples were prepared as a background signal control (n = 1) and negative spiking control (n = 1) during sample processing, respectively. Following spiking, tubes were placed on a ThermoScientific Solaris 2000 shaker at 150 rpm for 3 hours at 22°C. Next, samples were centrifuged at 20,000 *g* for 20 min at 4°C. After, 200 µL of supernatant was collected for DNA extraction, and the remaining liquid was placed into a new 50 mL conical tube and stored at 4°C. Approximately 250 mg of dewatered solids was retrieved and used for DNA extraction. If 250 mg of solids were not available, all solids from that sample were used. The mass of each sample was taken and recorded for future nucleic acid quantification, following a previously reported methodology (Roldan-Hernandez et al., 2023). Any remaining solids were placed into 2 mL tubes and stored at 4°C. Samples were spiked and processed within 48 hours of collection, then underwent DNA extraction using the Qiagen AllPrep PowerViral kit. An extraction blank was included as a negative process control (n = 1). Sample extracts were stored at -80°C until assayed within 1 month of extraction. For the dPCR of 10^4^ liquids samples, four wells were dedicated to each sample, analyzed using the HyperWell setting within the Qiacuity software and accounted for in the dimensional analysis. All other samples were quantified using single wells.

### 2.4. Viral nucleic acid quantification

dPCR was used to quantify all viral nucleic acids in this study on a Qiacuity dPCR platform. VV and HI-MPXV were quantified in samples using the singleplex non-variola orthopoxvirus assay (NVAR). Table 1 shows the characteristics of each assay used. Each plate included two wells for no template controls (NTCs) and two wells of a positive control plasmid (cat. no. RGTM 10223, NIST). Three positive partitions per sample were required for the sample to be considered positive. Additional details on dPCR reactions can be found in the Supplemental Materials.

**Table 1:**
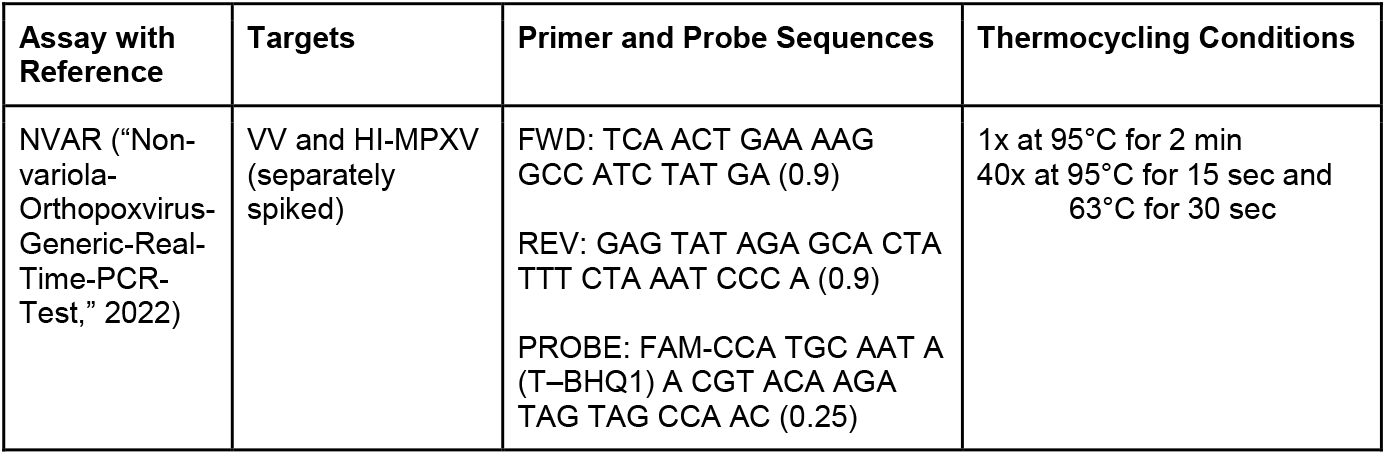
ddPCR primers, probe, and conditions for assay used in this study. Concentrations of primers and probes (in µM) in each reaction are shown in parentheses following sequences.

### 2.5. Data Analysis

Data generated by dPCR in gc/µL of bulk reaction was exported from the Qiacuity platform program. Dimensional analysis was used to convert sample concentrations from gc/µL to gc/mL for primary influent and DNAse/RNAse-free water samples and gc/g of dry solids for settled solids samples. More details on the data analysis, including dimensional analysis equations, level of detection for dPCR, and gene copies per positive partition, are in the SM. The threshold for positive partitions was set individually for each dPCR cassette and the same threshold was used for all wells within a plate. Data was viewed under the “Histogram” tab of the Qiacuity software to identify the lowest and highest fluorescence peaks. The average of these two peaks was taken, and this value was used to set the threshold.

For the partitioning experiment, the linear Freundlich isotherm model was investigated here, as this adsorption model has been commonly used in previous wastewater virus partitioning studies(Yang et al., 2022; Roldan-Hernandez and Boehm, 2023; Roldan-Hernandez et al., 2024). The following linear equation was used for the Freundlich isotherm model:

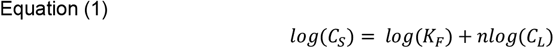

where C_S_ is the equilibrium concentration of gene copies in solids (gc/g dry solids), K_F_ is the Freundlich adsorption capacity, n is the adsorption intensity, and C_L_ is the equilibrium concentration of gene copies in the liquid portion of influent.

Microsoft Excel (version 16.16.27) was used to record all data. R (version 4.4.1) and RStudio (version 2024.04.2+764) were used for data analysis. The lm and summary functions were used to run the linear regression models, report r^2^ values, and calculate standard errors (SE). GraphPad Prism (version 10.3.1) was used to visualize figures, as well as to conduct significance testing. A paired, two-tailed t-test was used to assess the significance of virus type on decay rates, while a two-way ANOVA with Tukey’s multiple comparisons tests was used to assess the significance of matrices and temperatures on decay rates. The level of significance was set to a value of α = 0.05 for both tests. A Shapiro-Wilk test was also run to ensure normality of the dataset prior to significance testing; the resulting QQ plots from those tests are provided in Figure S1.

## 3. Results

### 3.1 Quality Assurance and Control

At least 3 positive partitions were required for a sample to be considered above the theoretical limit of detection, with that being 0.3 gene copies per microliter (gc/µL) of template for a single well. When combining same-sample wells using the HyperWell setting, the theoretical limit of detection for four wells was 0.075 gc/µL of template and 0.05 gc/µL of template for six wells. The analytical limit of detection was determined to be 0.205 gc/µL of template, which was calculated following previously outlined methods (Armbruster and Pry, 2008) and recommended by the Qiagen dPCR Handbook (“Qiagen Microbial DNA dPCR Handbook,” 2022). For all experiments, non-template controls (NTCs) for dPCR and extraction blanks were below the limit of detection. For the methods evaluation experiment, all primary influent no-spike samples were below the limit of detection, while one settled solids no-spike sample was at the theoretical limit of detection (3 positive partitions). For the persistence experiment, all no-spike controls were below the limit of detection. For the partitioning experiment, no-spike controls for the liquids fraction were below the limit of detection, while the solids fraction had an average concentration of 5800 gc/g. The NTCs had an average of 25,321 valid partitions (SE = 757.8; 95% CI ± 171.5). The NVAR plasmid used as a positive control for dPCR had an average reaction concentration of 3,006.4 gc/µL (SE = 177.2 gc/µL; 95% CI ± 54.2 gc/µL), 24,592 valid partitions (SE = 1483.7; 95% CI ± 454.1), and 22,047 positive partitions (SE = 1432.8; 95% CI ± 438.6).

### 3.2. Methods Evaluation

As wastewater is a complex matrix to quantify viral genetic markers (Kantor et al., 2021), we first compared several combinations of frequently used concentration and extraction methods to determine trends in recovery of HI-MPXV and VV DNA markers in two commonly used matrices: primary settled solids and primary influent. Although we spiked HI-MPXV and VV to greater than 10^4^ gc/g of wet solids, we saw low recovery of the poxviruses from primary settled solids. Percent recovery ranged from 0.01% to 1.9% for concentrated influent samples, while unconcentrated molecular process controls (MPCs) had higher recoveries, from 4.5% to 41%. Percent recoveries were in general higher for primary influent than for settled solids, which is consistent with previous comparisons of primary influent and settled solids methodologies (Table 2).

**Table 2:**
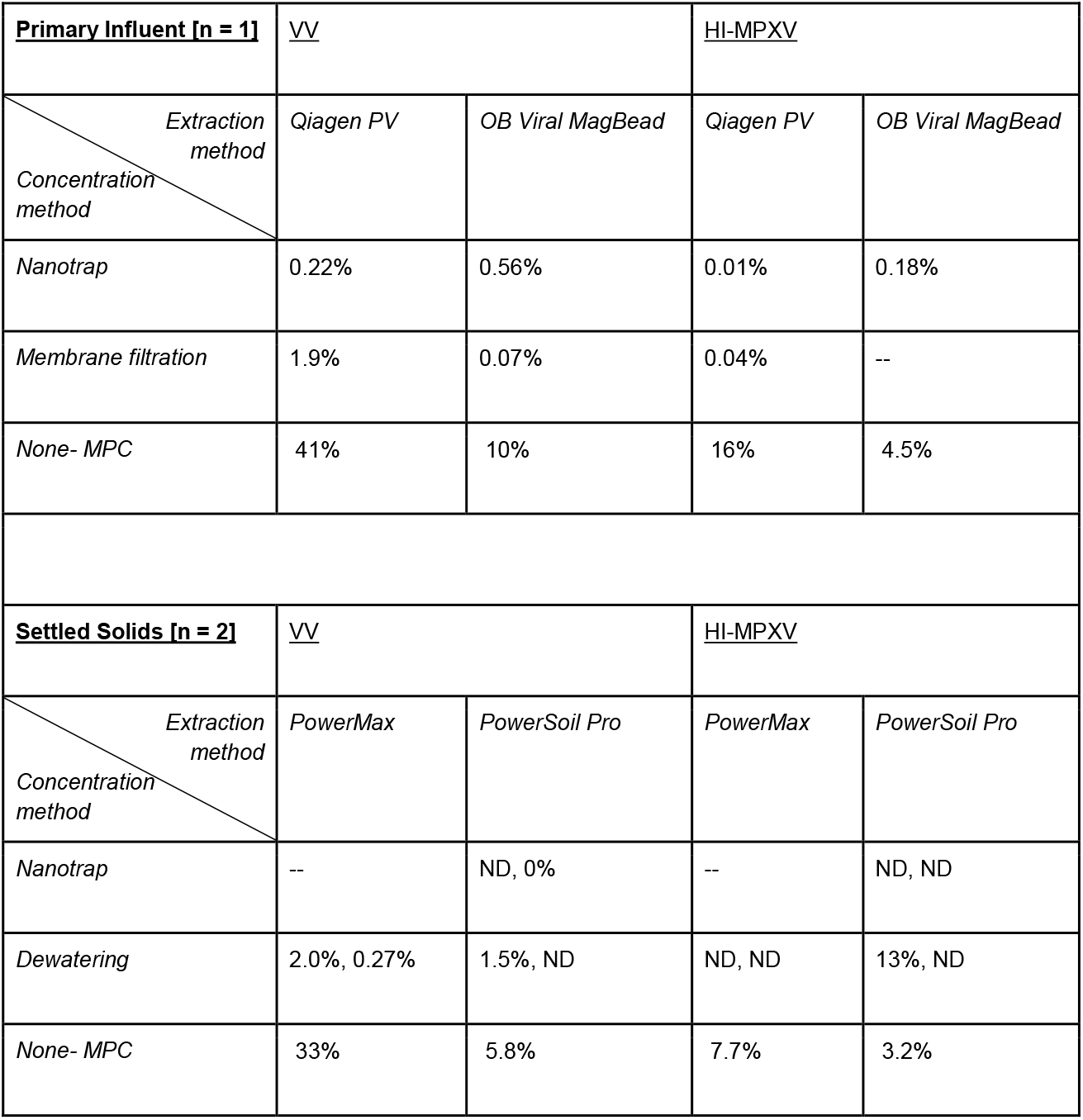
Average percent recovery of HI-MPXV and VV in primary influent (primary influent, n = 1 site) and primary settled solids (primary settled solids, n = 2 sites) after spiking. MPC = molecular process control: a primary influent and settled solids sample that was not concentrated and spiked with virus stock. Dashes indicate that a given combination was not tested.

| **Primary Influent [n = 1]** |  | <u>VV</u> |  | <u>HI-MPXV</u> |  |
| --- | --- | --- | --- | --- | --- |
| *Extraction*<br>*method*<br>*Concentration*<br>*method* |  | *Qiagen PV* | *OB Viral MagBead* | *Qiagen PV* | *OB Viral MagBead* |
| *Nanotrap* |  | 0.22% | 0.56% | 0.01% | 0.18% |
| *Membrane filtration* |  | 1.9% | 0.07% | 0.04% | -- |
| *None- MPC* |  | 41% | 10% | 16% | 4.5% |
| **Settled Solids [n = 2]** |  | <u>VV</u> |  | <u>HI-MPXV</u> |  |
| *Extraction*<br>*method*<br>*Concentration*<br>*method* |  | *PowerMax* | *PowerSoil Pro* | *PowerMax* | *PowerSoil Pro* |
| *Nanotrap* |  | -- | ND, 0% | -- | ND, ND |
| *Dewatering* |  | 2.0%, 0.27% | 1.5%, ND | ND, ND | 13%, ND |
| *None- MPC* |  | 33% | 5.8% | 7.7% | 3.2% |

\* Qiagen PV = Qiagen AllPrep PowerViral DNA/RNA kit; OB Viral MagBead = Omega Biotek MagBead Viral DNA/RNA kit; PowerMax = Qiagen DNeasy PowerMax Soil kit; PowerSoil Pro = Qiagen PowerSoil Pro kit

Due to higher recoveries in influent samples without concentration compared to those with concentration steps added, we moved forward without sample concentration and with the Qiagen AllPrep PowerViral DNA/RNA kit for primary influent. For primary settled solids samples, each extraction kit had relatively similar performance so we continued with the Qiagen PowerSoil Pro kit as it also had a faster processing time and performed better than the PowerMax kit when recovering HI-MPXV from dewatered settled solids samples (Table 1). We also performed an inhibition assessment for each primary influent and settled solids sample by diluting 1:10 prior to dPCR quantification. We found <10% inhibition for all but two samples, those being the HI-MPXV and VV dewatered primary settled solids samples extracted using the PowerSoil Pro kit. However, since little to no inhibition was evident in most samples, we decided not to continue with dilutions for the persistence and partitioning experiments.

### 3.3. Persistence of HI-MPXV and VV

Chick’s Law was used to investigate differences in decay rates between viruses, matrices, and temperatures, which are shown in Figure 1. Linear first-order decay was found to be a good fit for all influent samples, with r^2^ ranging from 0.890 to 0.959. In settled solids and DNAse/RNAse-free water, r^2^ were lower, ranging from 0.052 to 0.680 and 0.060 to 0.646, respectively. Decay rates (k) and their SE, r^2^, RMSE, and T_90_ values are found in Table 3.

**Table 3:**
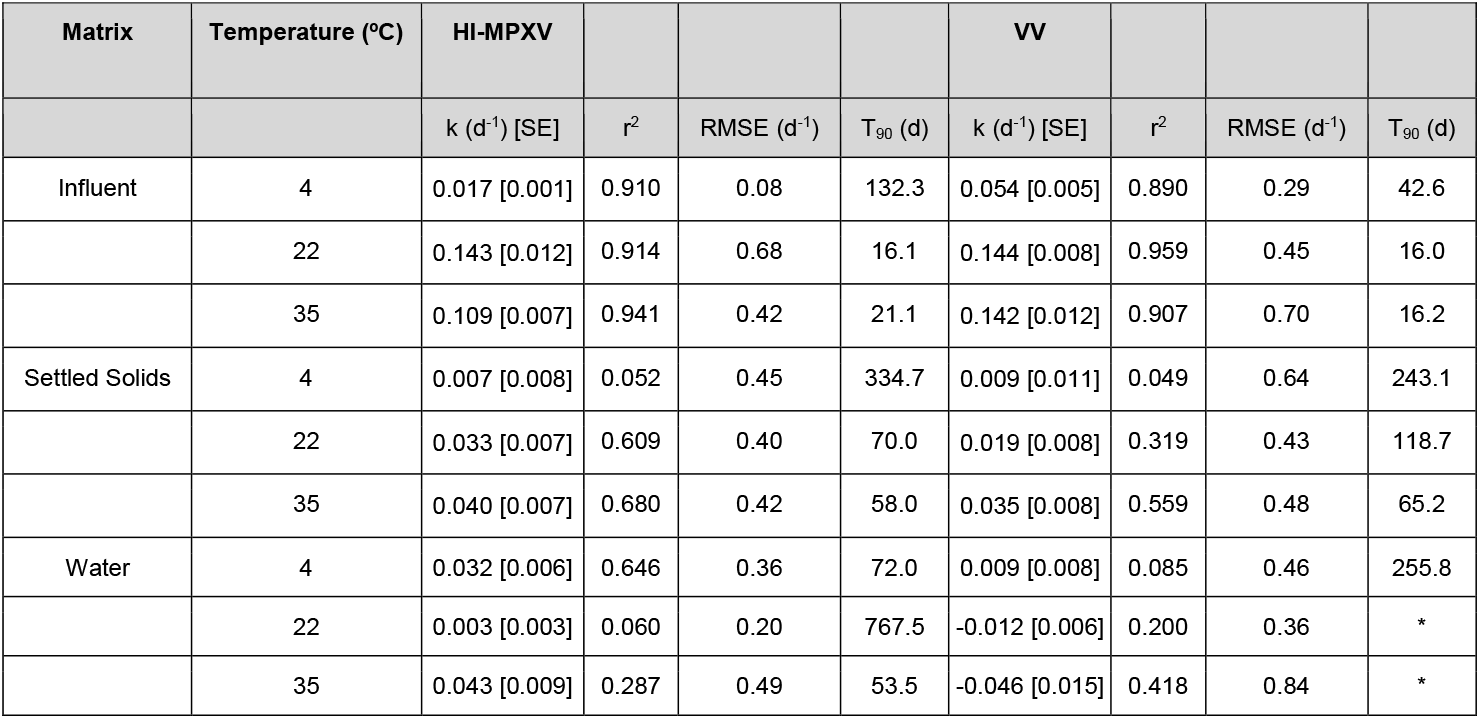
Decay rate constants (k) for HI-MPXV and VV in primary influent, primary settled solids, and DNase/RNase-free water mesocosms. Each mesocosm (in biological triplicates) was stored at each temperature in the dark and subsampled over 30 days. Chick’s Law was used to generate decay rate constants, r^2^, RMSE, and T_90_ values after confirming reasonable fit (r^2^ > 0.85). Asterisks for water at 22°C and 35°C indicate that T_90_ values were not calculated due to a negative decay rate constant.

| Matrix | Temperature (°C) | HI-MPXV |  |  |  | VV |  |  |  |
| --- | --- | --- | --- | --- | --- | --- | --- | --- | --- |
| | | k (d<sup>-1</sup>) [SE] | $r^2$ | RMSE (d<sup>-1</sup>) | $T_{90}$ (d) | k (d<sup>-1</sup>) [SE] | $r^2$ | RMSE (d<sup>-1</sup>) | $T_{90}$ (d) |
| Influent | 4 | 0.017 [0.001] | 0.910 | 0.08 | 132.3 | 0.054 [0.005] | 0.890 | 0.29 | 42.6 |
|  | 22 | 0.143 [0.012] | 0.914 | 0.68 | 16.1 | 0.144 [0.008] | 0.959 | 0.45 | 16.0 |
|  | 35 | 0.109 [0.007] | 0.941 | 0.42 | 21.1 | 0.142 [0.012] | 0.907 | 0.70 | 16.2 |
| Settled Solids | 4 | 0.007 [0.008] | 0.052 | 0.45 | 334.7 | 0.009 [0.011] | 0.049 | 0.64 | 243.1 |
|  | 22 | 0.033 [0.007] | 0.609 | 0.40 | 70.0 | 0.019 [0.008] | 0.319 | 0.43 | 118.7 |
|  | 35 | 0.040 [0.007] | 0.680 | 0.42 | 58.0 | 0.035 [0.008] | 0.559 | 0.48 | 65.2 |
| Water | 4 | 0.032 [0.006] | 0.646 | 0.36 | 72.0 | 0.009 [0.008] | 0.085 | 0.46 | 255.8 |
|  | 22 | 0.003 [0.003] | 0.060 | 0.20 | 767.5 | -0.012 [0.006] | 0.200 | 0.36 | * |
|  | 35 | 0.043 [0.009] | 0.287 | 0.49 | 53.5 | -0.046 [0.015] | 0.418 | 0.84 | * |

**Figure 1:**
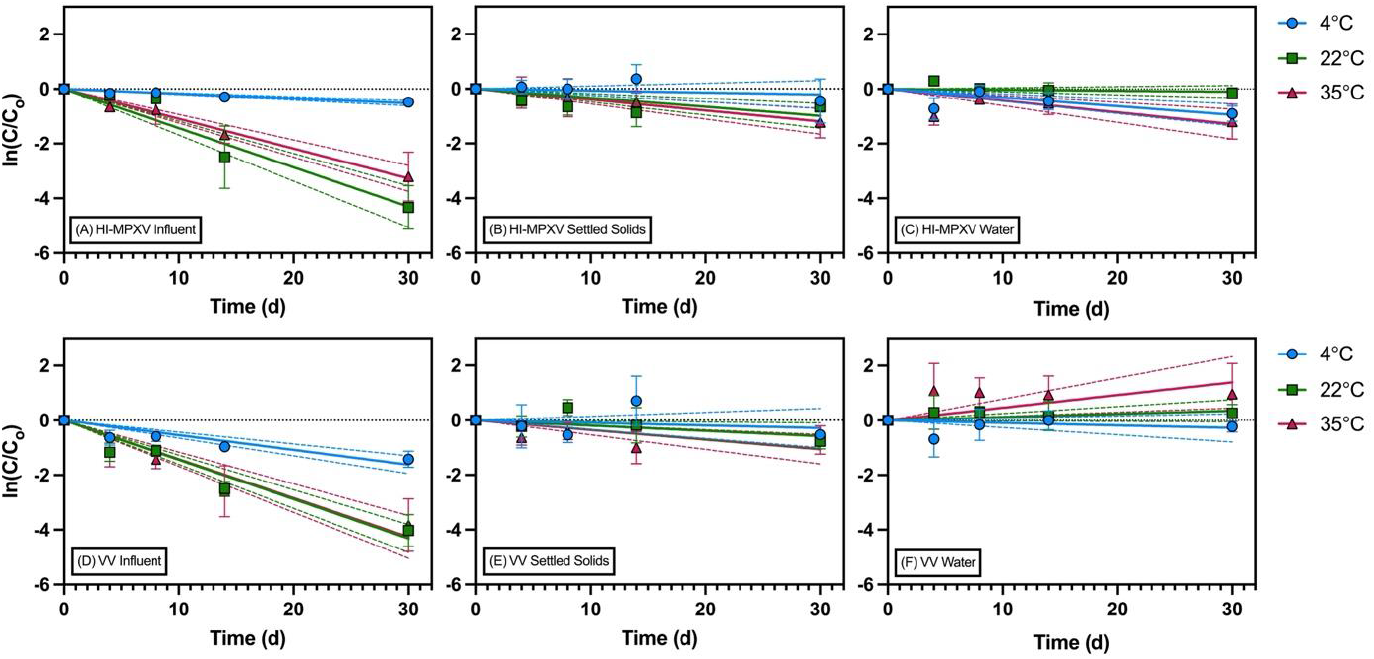
Persistence of HI-MPXV (top row) and VV (bottom row) over 30 days across primary influent, settled solids, and DNAse/RNAse-free water. Samples were held at 4°C (blue), 22°C (green), or 35°C (red) throughout the study. Dashed lines represent the 95% confidence intervals for the slope, while error bars indicate the standard deviation among mesocosm triplicates

The decay rate constants (k) for each virus were significantly lower at 4°C compared to 22°C or 35°C for primary influent (Tukey’s Multiple Comparisons test, p=0.0051 and p=0.0147, respectively). Little decay of both viruses was observed at 4°C in primary influent. In settled solids, the k-values between different temperatures were not significantly different. Finally, in DNAse/RNAse-free water, we also found the k-values to not be significantly different between temperatures. A complete list of multiple comparisons and p-values from our Tukey’s Multiple Comparisons test can be found in Table S1. As shown in Figure 2, when compared to the decay rates of RNA viruses from previously published studies, HI-MPXV and VV exhibited very similar decay to other viruses at their respective temperatures, including SARS-CoV-2 and Zika virus (ZIKV) in primary influent and SARS-CoV-2 in settled solids. A full list of study names, decay rates, and T_90_ values used in the creation of Figure 2 can be found in Table S2 for primary influent and Table S3 for settled solids.

**Figure 2:**
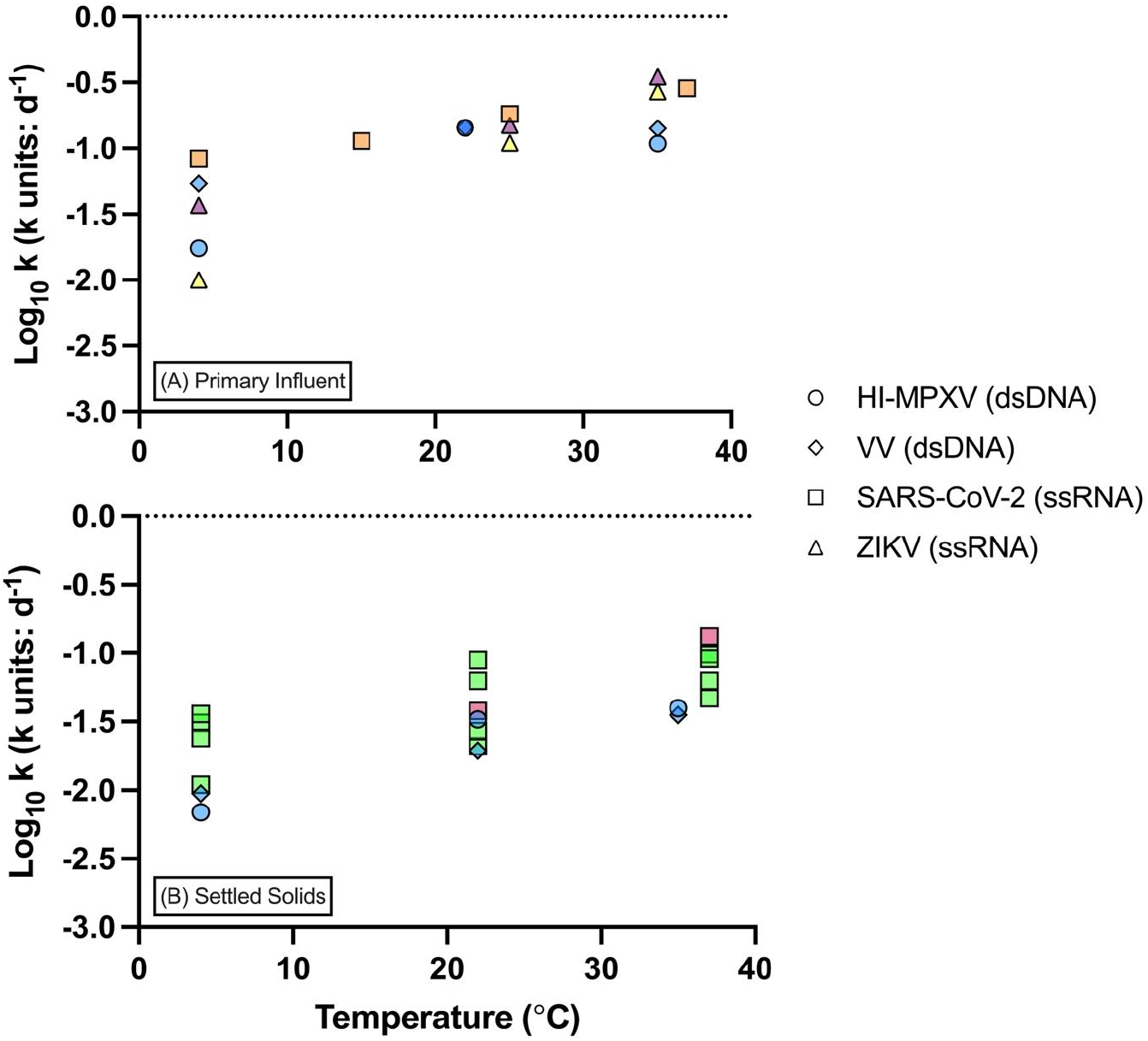
Comparison of log_10_ k decay values from this study with those of other published RNA viruses. All data was generated using molecular methods as opposed to culture-based assays. Circles represent HI-MPXV (our study), diamonds represent VV (our study), squares represent SARS-CoV-2 (Ahmed et al., 2020a; Roldan-Hernandez et al., 2022; Zhang et al., 2024), and triangles represent ZIKV (Muirhead et al., 2020; Zhu et al., 2023). Colors denote different papers (n = 5), with specific k-values and study names listed in Table S1 for primary influent and Table S2 for settled solids.

### 3.4. Solid-Liquid Partitioning

Using digital PCR, the log_10_-transformed virus concentrations were calculated for the liquids (C_L_, gc/mL) and solids (C_S_, gc/g). For HI-MPXV, C_L_ ranged from 420 to 615,600 gc/mL and C_S_ ranged from 1.99 x 10^6^ to 4.01 x 10^8^ gc/g. For VV, C_L_ ranged from 300 to 37,500 gc/mL and C_S_ ranged from 2.27 x 10^5^ to 2.83 x 10^7^ gc/g. Samples originally spiked at a virus concentration of 10^3^ gc/mL had no liquid samples above the limit of detection, but viral DNA markers were found in the solids fraction (X’s on y-axis in Figure 3). The adsorption capacity, K_F_, was calculated by exponentiating the y-intercept of the linearized isotherms. The K_F_ for HI-MPXV was 31,800 mL/g, with SE bounds of 18,600 to 54,600 mL/g. For VV, the K_F_ was 1,000 mL/g, with SE bounds of 700 to 1,400 mL/g. The adsorption intensity, n, was determined by taking the reciprocal of the isotherm slopes. The slopes were found to be significantly different (two-tailed test, p=0.0054) between the two viruses. For HI-MPXV, the n was found to be 1.41 (SE ± 0.10), while n for VV was 1.01 (SE ± 0.04). The r^2^ was calculated to be 0.978 for HI-MPXV and 0.994 for VV.

**Figure 3:**
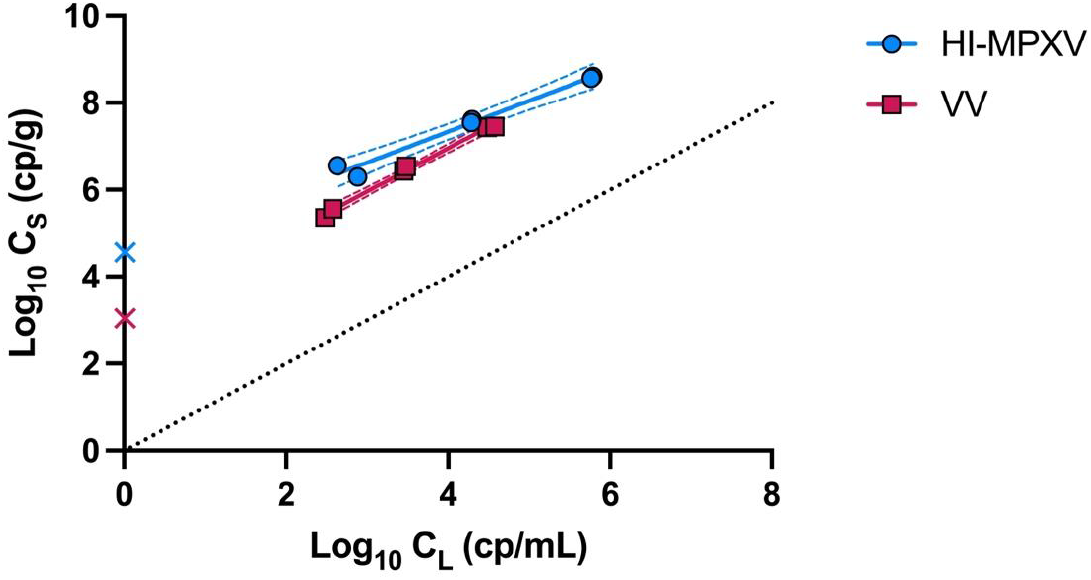
Freundlich adsorption isotherms for the solid-liquid partitioning of HI-MPXV (blue) and VV (red) at different spike concentrations. X’s on the y-axis indicate that for the primary influent sample spiked with 10^3^ gc, there was only amplification of the DNA marker in the solids fraction and not in the liquid fraction (BLD); these values were excluded from the linear regression as there was no amplification in the liquids. Dashed lines represent a 95% confidence interval for each regression. A dotted line of identity (y = x) is provided.

## 4. Discussion

There is sparse data concerning the decay and partitioning behaviors of orthopoxviruses across different wastewater matrices, despite their public health importance and the documented use of wastewater surveillance to assist with the identification of outbreaks. Likewise, it is imperative to determine the optimal and appropriate methodologies regarding the recovery of viral nucleic acids from a wastewater sample (Li et al., 2024). In this study, we provide quantitative information on the behavior of two representative orthopoxviruses to guide future surveillance of orthopoxviruses via wastewater, in particular, related to appropriate storage conditions, decay rates, and adsorption parameters that are important for wastewater-based modeling of cases.

First, we found that the recovery of viral markers was greater in primary influent compared to settled solids, and we observed substantial losses during the use of concentration methods for both matrices. Higher recoveries were obtained for VV than for HI-MPXV from both matrices; since we spiked VV at an order of magnitude higher than HI-MPXV, the starting concentration may confound this assessment. Nevertheless, our results provided methodological information that we used to evaluate the persistence and partitioning characteristics of orthopoxvirus DNA markers in the next experiments.

In our persistence experiments, we found that the DNA markers for our orthopoxvirus surrogates were extremely persistent, especially in settled solids. The highest decay rates were observed in primary influent; both viruses decayed significantly faster compared to settled solids at 22°C and 35°C, as well as water at 22°C and 35°C. Likewise, temperature had a significant impact on decay for both viruses at 4°C compared to 22°C and 35°C in primary influent. Once more, p-values for these comparisons are found in Table S1. Interestingly, for HI-MPXV, the decay regressions are very similar between settled solids and nuclease-free water, signifying the solids’ ability to provide a certain level of protection from microbial predation and/or enzymatic reactions that degrade viral particles. This indicates that specific storage conditions of settled solids may allow for better retroactive assessment of non-variola orthopoxviruses compared to primary influent and may be preferable for wastewater-based epidemiology practices and monitoring. Additionally, the decay rates of both viruses, HI-MPXV and VV, were not significantly different (paired t-test, p=0.5258), implying that the slight differences between the two orthopoxviruses do not have a meaningful effect on their decay rates in our experiments. We found that matrix significantly affected decay (Two-Way ANOVA, p=0.0002), as did the interaction between matrix and temperature (Two-Way ANOVA, p=0.034), accounting for 60.32% and 20.39% of variation, respectively. The T_90_ time for raw primary settled solids was 2.5-4.3 times longer for HI-MPXV and 4.0-7.4 times longer for VV in comparison to primary influent, indicating that sampling programs that collect a variety of wastewater sample types could leverage primary settled solids samples for retrospective analysis of orthopoxvirus DNA marker dynamics.

In general, DNA markers have been considered to be more persistent than RNA markers in environmental samples (Inoue et al., 2023; Lobos et al., 2024). To estimate if this was true for the viral DNA markers in this study, we compared our values to previously estimated decay rates for RNA viruses calculated in primary influent and settled solids matrices (Figure 2). We found that the DNA-marker decay rate constants generated in this study surprisingly appeared to be very comparable to previously reported viral RNA marker decay rates, indicating that many RNA virus markers are not vastly less persistent than DNA virus markers, though some sample-to-sample variation may affect these specific ranges.

The fate of orthopoxviruses in wastewater has critical implications for wastewater sampling designs; for instance, the adsorption behavior of viral markers may impact what type of sample should be collected. The viral DNA markers for both orthopoxviruses were heavily sorbed to the solid fraction of primary influent. Even when primary influent was spiked with 10^3^ gc/mL of each virus during partitioning experiments, viral DNA markers were only recovered in the solid fraction, while the markers were below the LoD in the liquid fraction. This suggests that a concentration of greater than 10^3^ gc/mL is necessary in raw primarily influent for the viable quantification of the DNA marker in the clarified liquid fraction of influent samples, which will be a useful metric for practitioners recovering orthopoxviruses for wastewater-based epidemiology applications.

The adsorption behavior of the enveloped viruses in this study are consistent with other studies on enveloped viruses in wastewater. In a previous study (Roldan-Hernandez and Boehm, 2023), the Freundlich adsorption model parameters of enveloped viruses SARS-CoV-2 and RSV-A were found to have K_F_ values of 18,000 and 32,000 mL/g and n values of 0.81 and 1.24, respectively. Following that (Roldan-Hernandez et al., 2024), a follow-up study found a range of K_F_ values from 500 to 7,600 mL/g and n values from 0.93 to 1.14 for various enveloped viruses, including Dengue, West Nile, Zika, Influenza A, and SARS-CoV-2. We found the K_F_ values of HI-MPXV and VV, also enveloped viruses, to be 31,800 mL/g and 1,000 mL/g, respectively, and significantly different. These values mirror some of both the highest and lowest K_F_ values for enveloped viruses reported in the literature. Likewise, our n values for HI-MPXV and VV were 1.41 and 1.01, respectively. In particular, the adsorption intensity of HI-MPXV was very high, even exceeding those in the aforementioned investigations. Given the high similarity in genome size and virion diameter between HI-MPXV and VV (Johnson et al., 2006; Lu et al., 2023) as well as near-identical capsid compositions (Su et al., 2005), it seems unlikely, but possible, that the differences in adsorption capacity and intensity are driven by capsid protein binding parameters alone. However, the heat inactivation of the HI-MPXV used in this study may have driven some level of capsid protein degradation that differentiated its partitioning to solids, relative to the non-heat inactivated VV.

## 5. Limitations, Conclusions, and Future Work

While this study is comprehensive in its scope and design, we did not include infectivity assessment for HI-MPXV or VV, as our goal was to inform WBE practices that rely on molecular measurements of DNA and RNA markers. However, a recent study investigated the stability of mpox in wastewater over a 20-day time period at 16°C (Yinda et al., 2023). They utilized a culture-based assay to track mpox persistence in irradiated wastewater, and their data showed evidence of biphasic decay. We extracted their data and determined the k-values for each phase, assuming the first phase to be days 0 to 15 and the second phase to be days 15 to 20. We calculated decay rates of 0.033/day and 0.057/day for the first and second phases, respectively. These k-values are similar to the decay constants we found in our study for settled solids samples at 22°C (0.033/day) and 35°C (0.040/day). Additionally, the spike-in strains that we used in our study are not environmental isolates, which can be more persistent than laboratory strains (Ogbunugafor et al., 2010).

Nevertheless, our results have implications for environmental monitoring programs and retrospective analysis of archived wastewater samples. The findings of this work provide critical insight into wastewater sample collection, storage, and processing for agencies aiming to investigate community health and the prevalence of mpox cases within a sewershed. Primary settled solids samples exhibited the least decay at 4°C over the 30-day study period, potentially making them more suitable than primary influent for retrospective analyses of mpox community infection. Furthermore, decay rates and partitioning parameters were empirically determined and are comparable to other viruses investigated in similar studies, indicating that overall, primary settled solids stored at 4°C can be used to effectively identify viral DNA markers over a timeframe of up to 30 days (<1 log-decay for both HI-MPXV and VV), without any adjuvants needed. We included both HI-MPXV and VV in this investigation to better understand the generalizability of WBE parameters among poxviruses by comparing the two, providing insight into the behavior of dsDNA viruses in primary influent and settled solids, which we found sparse data on prior to this study. Over the course of our research, we have identified areas of future work to further understanding of decay kinetics and partitioning characteristics. For example, we identified very few recent studies exploring these characteristics for other pathogenic DNA viruses, such as *Herpesviridae* and *Papillomaviridae*, across various wastewater matrices. Moreover, the significant (but small) differences in adsorption parameters between HI-MPXV and attenuated VV observed in our work warrants further inquest into how viral treatments for laboratory research, such as chemical treatments or heat inactivation that decrease the biosafety level restrictions to work with biohazardous samples, affect adsorption behaviors.

## CRediT Authorship Contribution Statement

**Jacob R. Phaneuf:** Writing – Review and Editing, Writing – Original Draft, Data Curation, Visualization, Methodology, Investigation, Formal Analysis. **Gyuhyon Cha:** Conceptualization, Writing – Review and Editing. **Janet K. Hatt:** Writing – Review and Editing. **Konstantinos T. Konstantinidis:** Resources, Writing – Review and Editing. **Katherine E. Graham:** Writing – Review and Editing, Writing – Original Draft, Data Curation, Methodology, Investigation, Formal Analysis, Funding Acquisition, Resources, Supervision, Conceptualization

## Data Availability

The raw data required to reproduce the above findings are available to download from the following link: https://hdl.handle.net/1853/76625

## Acknowledgments

We thank Kevin Zhu for his feedback on our experimental design. We thank the staff of POTW-A and POTW-B for providing wastewater samples and their corresponding physicochemical data. We thank the Georgia Institute of Technology for the funds used to complete this research. BioRender (https://BioRender.com) was used to create the graphical abstract.

