## Supplementary Material for "Decay and Solid-Liquid Partitioning of Mpox and Vaccinia Viruses in Primary Influent and Settled Solids to Guide Wastewater-Based Epidemiology Practices"

Number of Pages: 8

Number of Tables: 3

Number of Figures: 2

**Methods**

**Sample concentration:** The Nanotrap bead method proceeded as follows: 1.2 mL of primary settled solids was added to a tube with 10.8 mL of Nanotrap Buffer 2 and three glass grinding beads (approx. 0.5 cm diameter). Tubes were vortexed for 10 min at maximum speed, and afterwards solids were allowed to settle for 10 min at room temperature. Ten mL of the resulting supernatant was added to a new tube with 150 µL of Nanotrap Microbiome A particles, inverted to mix, and incubated for 10 min at room temperature with an inversion to mix halfway through the incubation period. Tubes were then placed on a magnetic rack for 5 min and the supernatant was removed. One mL of Nanotrap Buffer 2 was added to the resulting pellet and it was resuspended by pipetting and washing the sides of the tube, prior to pipetting into a new microcentrifuge tube. Tubes were then placed on a magnetic rack for 5 min and the resulting supernatant was removed by pipetting. Lysis buffer (500 µL) of lysis buffer was added to each pellet and pipetted up and down to resuspend, then heated at 95°C for 10 min on a heat block. Finally, heated suspensions were placed on a magnetic rack for 5 min and 400 µl of resulting supernatant was added to a new tube that was placed at 4°C until the remaining nucleic acid extraction steps could be completed (within 48 hours). The dewatering method started with centrifuging ~20 g aliquots of raw primary settled solids at 24,000xg for 30 min at 4°C, then removing the supernatant from each tube. The resulting dewatered solids were added to extraction tubes and weighed. After concentration, all concentrates were stored in lysis buffer at 4°C prior to remaining nucleic acid extraction steps.

In brief, the membrane filtration method started by measuring 10 mL of influent and spiking with a 10 M solution of MgCl_2_ to a final concentration of 0.05 M and inverting to mix. After 5 min, the solution was filtered through a magnetic filtration funnel with a 47 mm diameter membrane filter (0.45 µm pore size, 47 mm diameter, mixed-cellulose esters, gamma-irradiated). Membranes were removed from filtration funnels after all liquid had passed through and placed into microcentrifuge tubes containing the lysis buffer for each respective kit tested. For each virus spiked, one membrane was filtered using 48 mL, and the resulting membrane filter was split into approximate halves prior to adding to two separate microcentrifuge tubes, one for each extraction kit. For the Nanotrap concentration method, we followed the manufacturer’s protocol, corresponding to APP-088: Nanotrap Microbiome A 10 mL Manual Protocol from Ceres (“APP-088 R02 Manual Nanotrap Microbiome A Enviro Water Protocol using AllPrep PowerViral DNA RNA Mini,” 2024).

**Copies Per Partition for dPCR:** Copies per partition (λ) was determined using the following formula provided in the QIAcuity User Manual Extension (“QIAcuity Manual Extension,” 2021.):

$$\lambda=-ln(\frac{Number of Valid Partitions-Number of Positive Partitions}{Number of Valid Partitions})$$

The average $\lambda$ for both viruses across each matrix were found to be 0.0480 for primary influent, 0.0003 for settled solids, and 0.0237 for DNAse/RNAse-free water.

**Dimensional Analysis:** The following equations were used for dimensional analysis in the persistence and partitioning experiments:

Liquids:

$$Conc \left[ \frac{gc}{mL influent} \right]=dPCR Output \left[ \frac{gc}{{\mu L}_{rxn}} \right]\times dPCR Rxn \left[ \frac{{\mu L}_{rxn}}{{\mu L}_{template}} \right]\times Ext. Factor [\frac{{\mu L}_{elu}}{{\mu L}_{input}}]\times\frac{1000 \mu L}{1 mL}$$

Solids:

$$Conc\left[ \frac{gc}{g dry solids} \right]=dPCR Output \left[ \frac{gc}{{\mu L}_{rxn}} \right]\times dPCR Rxn \left[ \frac{{\mu L}_{rxn}}{{\mu L}_{template}} \right]\times Ext. Factor [\frac{{\mu L}_{elu}}{g dry solids}]$$

where “g dry solids” is the mass of the post-centrifuged solids (pellet only), “dPCR Output” is the raw concentration given by the QIAcuity system, “dPCR Rxn” is the volume per well in the QIAcuity nanoplate (40μL) over the volume of DNA extract added to the well (10μL), and “Ext. Factor” is the volume of DNA extract (100μL) obtained from DNA extraction over the volume or mass used as input for the extraction.

**Parameters for Two-Way ANOVA**: When using Prism to assess matrix/temperature significance, the following settings were applied: (1) from the “Model” tab, no factors were matched and a full model was fit and (2) from the “Multiple Comparisons” tab, cell means with others in rows and columns were used for the comparisons.

Table S1: Results of Tukey’s Multiple Comparisons Test performed in Prism to assess significance between sources of variation among mesocosms from different experimental conditions and the percent of total variation from matrix, temperature, and the interaction.

| ***Tukey's Multiple Comparisons Test*** | |  |  |
| --- | --- | --- | --- |
| **Influent** | **Significant?** | **p-value Summary** | **p-value** |
| 4C vs. 22C | Yes | ** | 0.0051 |
| 4C vs. 35C | Yes | * | 0.0147 |
| 22C vs. 35C | No | ns | 0.7602 |
| **Settled Solids** |  |  |  |
| 4C vs. 22C | No | ns | 0.7602 |
| 4C vs. 35C | No | ns | 0.4963 |
| 22C vs. 35C | No | ns | 0.8922 |
| **Water** |  |  |  |
| 4C vs. 22C | No | ns | 0.5979 |
| 4C vs. 35C | No | ns | 0.6680 |
| 22C vs. 35C | No | ns | 0.9922 |
| **4C** |  |  |  |
| Influent vs. Settled Solids | No | ns | 0.5407 |
| Influent vs. Water | No | ns | 0.8251 |
| Settled Solids vs. Water | No | ns | 0.8742 |
| **22C** |  |  |  |
| Influent vs. Settled Solids | Yes | ** | 0.0030 |
| Influent vs. Water | Yes | *** | 0.0006 |
| Settled Solids vs. Water | No | ns | 0.4748 |
| **35C** |  |  |  |
| Influent vs. Settled Solids | Yes | * | 0.0166 |
| Influent vs. Water | Yes | ** | 0.0018 |
| Settled Solids vs. Water | No | ns | 0.3136 |

##### Table S2: Comparison of k-values (d^-1^) and T_90_ values (d) in primary influent across cited literature using various viral markers. Colors correspond to data points found in Figure 2.

| **Study Name** | **Virus** | **Temperature [°C]** | **k [d-1]** | **T_90_ [d]** |
| --- | --- | --- | --- | --- |
| Our Study (Blue) | HI-MPXV | 4 | 0.017 | 132.3 |
|  |  | 22 | 0.143 | 16.1 |
|  |  | 35 | 0.109 | 21.1 |
|  | VV | 4 | 0.054 | 42.6 |
|  |  | 22 | 0.144 | 16.0 |
|  |  | 35 | 0.142 | 16.2 |
| "Decay of SARS-CoV-2 and surrogate murine hepatitis virus RNA in untreated wastewater to inform application in wastewater-based epidemiology" by Ahmed et al., 2020 (Orange) | SARS-CoV-2 | 4 | 0.084 | 27.8 |
|  |  | 15 | 0.114 | 20.4 |
|  |  | 25 | 0.183 | 12.6 |
|  |  | 37 | 0.286 | 8.0 |
| "Zika virus RNA persistence and recovery in water and wastewater: An approach for Zika virus surveillance in resource-constrained settings" by Zhu et al., 2023 (Magenta) | ZIKV | 4 | 0.037 | 62.2 |
|  |  | 25 | 0.150 | 15.4 |
|  |  | 35 | 0.350 | 6.6 |
| "Zika Virus RNA Persistence in Sewage" by Muirhead et al., 2020 (Yellow) | ZIKV | 4 | 0.010 | 230.3 |
|  |  | 25 | 0.110 | 20.9 |
|  |  | 35 | 0.270 | 8.5 |

##### Table S3: Comparison of k-values (d^-1^) and T_90_ values (d) in settled solids across cited literature using various viral markers. Colors correspond to data points found in Figure 2.

| **Study Name** | **Virus** | **Temperature [°C]** | **k [d-1]** | **T_90_ [d]** |
| --- | --- | --- | --- | --- |
| Our Study (Blue) | HI-MPXV | 4 | 0.007 | 334.7 |
|  |  | 22 | 0.033 | 70.0 |
|  |  | 35 | 0.040 | 58.0 |
|  | VV | 4 | 0.009 | 243.1 |
|  |  | 22 | 0.019 | 118.7 |
|  |  | 35 | 0.035 | 65.2 |
| "Persistence of Endogenous SARS-CoV‑2 and Pepper Mild Mottle Virus RNA in Wastewater-Settled Solids" by Roldan-Hernandez et al., 2022 (Green) | SARS-CoV-2 (N1, POTW-A) | 4 | 0.024 | 95.0 |
|  |  | 22 | 0.027 | 85.9 |
|  |  | 37 | 0.063 | 36.6 |
|  | SARS-CoV-2 (N1, POTW-B) | 4 | 0.036 | 64.6 |
|  |  | 22 | 0.063 | 36.5 |
|  |  | 37 | 0.091 | 25.3 |
|  | SARS-CoV-2 (N2, POTW-A) | 4 | 0.011 | 214.7 |
|  |  | 22 | 0.021 | 107.3 |
|  |  | 37 | 0.047 | 49.4 |
|  | SARS-CoV-2 (N2, POTW-B) | 4 | 0.031 | 75.4 |
|  |  | 22 | 0.089 | 26.0 |
|  |  | 37 | 0.098 | 23.5 |
| "Persistence of human respiratory viral RNA in wastewater-settled solids" by Zhang et al., 2024 (Red) | SARS-CoV-2 | 22 | 0.038 | 60.6 |
|  |  | 35 | 0.133 | 17.3 |

##### Figure S1: The QQ plots generated for persistence samples organized by virus and matrix following normality testing using the Shapiro-Wilk test in Prism. Temperature is denoted by different shapes and colors.

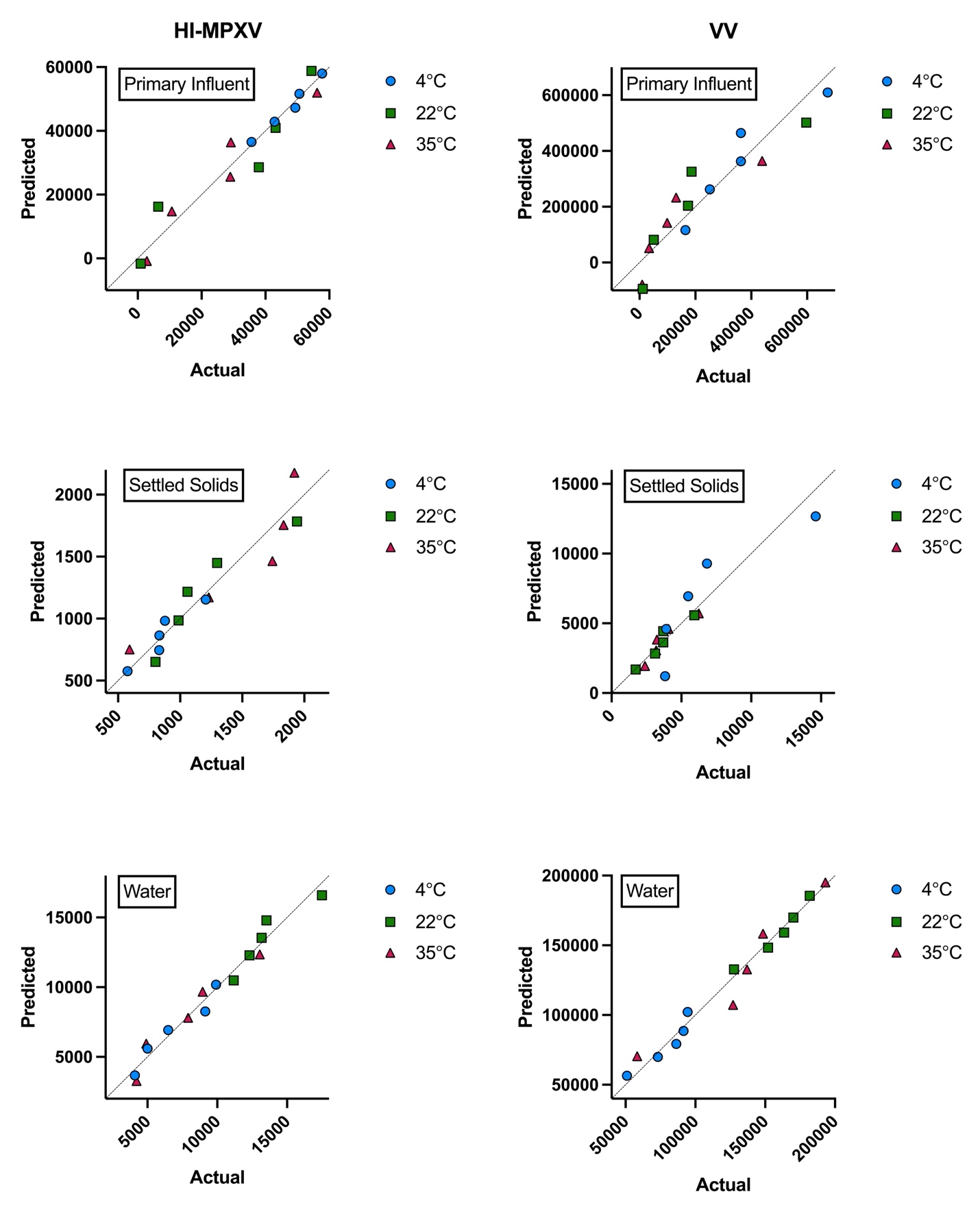

##### Figure S2: Environmental Microbiology Minimum Information (EMMI) checklist for reporting study details, control information, and experimental processes (Borchardt et al., 2021).

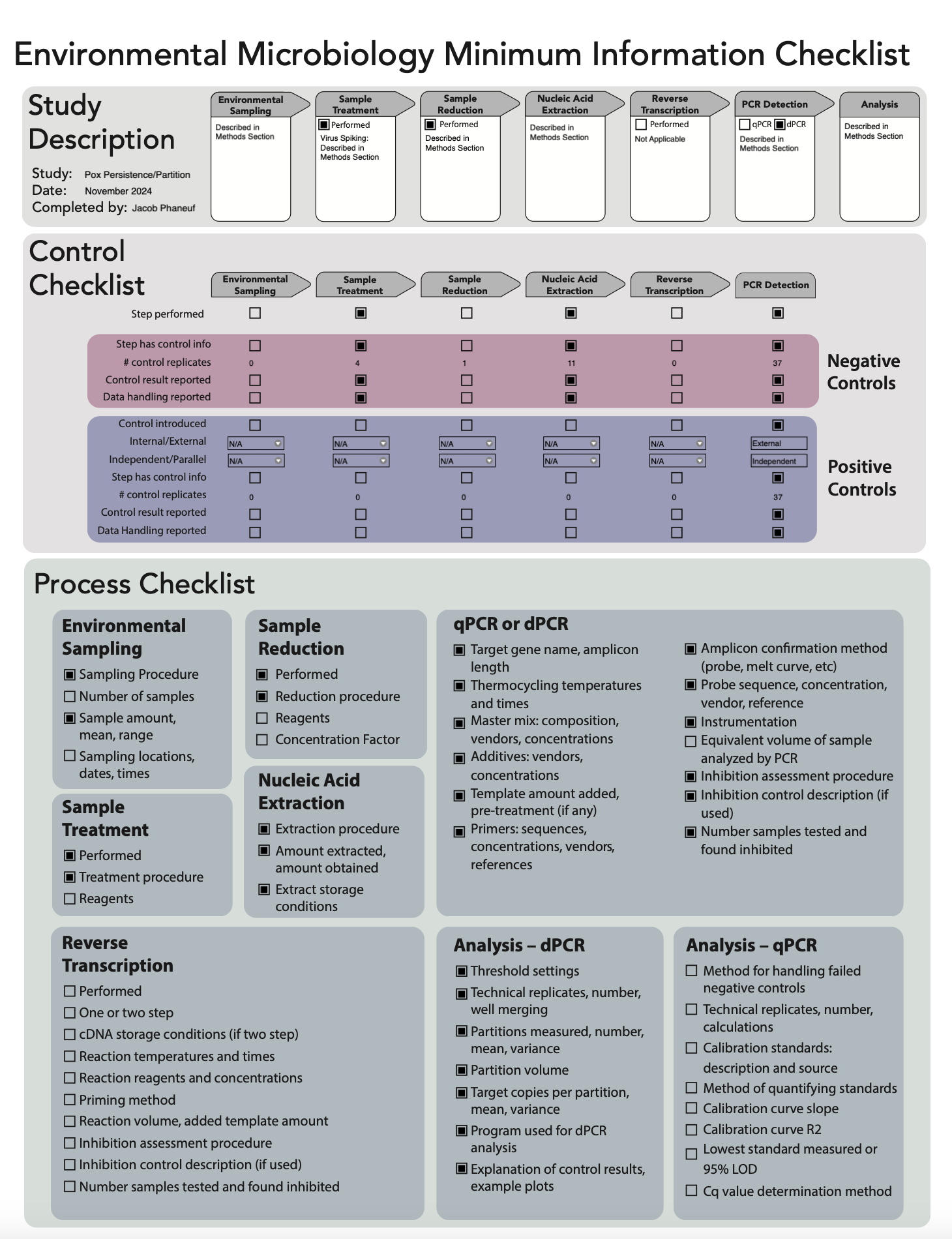
